# Inhibition of microbial biofuel production in drought stressed switchgrass hydrolysate

**DOI:** 10.1101/083758

**Authors:** Rebecca Garlock Ong, Alan Higbee, Scott Bottoms, Quinn Dickinson, Dan Xie, Scott A. Smith, Jose Serate, Edward Pohlmann, Arthur Daniel Jones, Joshua J. Coon, Trey K. Sato, Gregg R. Sanford, Dustin Eilert, Lawrence G. Oates, Jeff S. Piotrowski, Donna M. Bates, David Cavalier, Yaoping Zhang

## Abstract

**Background:** Interannual variability in precipitation, particularly drought, can affect lignocellulosic crop biomass yields and composition, and is expected to increase biofuel yield variability. However, the effect of precipitation on downstream fermentation processes has never been directly characterized. In order to investigate the impact of interannual climate variability on biofuel production, corn stover and switchgrass were collected during three years with significantly different precipitation profiles, representing a major drought year (2012) and two years with average precipitation for the entire season (2010 and 2013). All feedstocks were AFEX (ammonia fiber expansion)-pretreated, enzymatically hydrolyzed, and the hydrolysates separately fermented using xylose-utilizing strains of *Saccharomyces cerevisiae* and *Zymomonas mobilis.* A chemical genomics approach was also used to evaluate the growth of yeast mutants in the hydrolysates.

**Results:** While most corn stover and switchgrass hydrolysates were readily fermented, growth of S. *cerevisiae* was completely inhibited in hydrolysate generated from drought stressed switchgrass. Based on chemical genomics analysis, yeast strains deficient in genes related to protein trafficking within the cell were significantly more resistant to the drought year switchgrass hydrolysate. Detailed biomass and hydrolysate characterization revealed that switchgrass accumulated greater concentrations of soluble sugars in response to the drought and these sugars were subsequently degraded to pyrazines and imidazoles during ammonia-based pretreatment. When added *ex situ* to normal switchgrass hydrolysate, imidazoles and pyrazines caused anaerobic growth inhibition of *S. cerevisiae*.

**Conclusions:** In response to the osmotic pressures experienced during drought stress, plants accumulate soluble sugars that are susceptible to degradation during chemical pretreatments. For ammonia-based pretreatment these sugars degrade to imidazoles and pyrazines. These compounds contribute to *S. cerevisiae* growth inhibition in drought year switchgrass hydrolysate. This work discovered that variation in environmental conditions during the growth of bioenergy crops could have significant detrimental effects on fermentation organisms during biofuel production. These findings are relevant to regions where climate change is predicted to cause an increased incidence of drought and to marginal lands with poor water holding capacity, where fluctuations in soil moisture may trigger frequent drought stress response in lignocellulosic feedstocks.

## Background

Biofuels generated from lignocellulosic materials have enormous potential to reduce transportation-generated greenhouse gas emissions [1]. By 2030, the U.S. could be capable of supplying as much as 1.2 billion dry tons of agricultural residues and dedicated herbaceous energy feedstocks, enough to generate 58 billion gallons of ethanol per year [2]. However, biomass production in any given year is highly dependent on weather conditions. Soil moisture levels during a growing season are affected by both past and current levels of precipitation, and are a major determinant of lignocellulosic biomass yields in non-irrigated systems [3, 4]. Low levels of precipitation and soil moisture are particularly detrimental. Plants grown under water stressed conditions have reduced photosynthesis and slower growth, which reduces biomass yields [5, 6, 4]. Drought stress can also affect plant chemical composition, often resulting in reduced levels of structural carbohydrates [7–9] and accumulation of compounds that protect against osmotic stresses, including soluble sugars and amino acids (*e.g.*, proline) [5, 6]. These changes in plant composition are also predicted to result in lower ethanol yields from drought-stressed feedstocks [7, 8], although actual fermentations have never been carried out.

A number of different potential lignocellulosic bioenergy feedstocks are being considered in the U.S., including agricultural residues such as corn stover (*Zea mays* L.), and dedicated energy crops such as switchgrass (*Panicum virgatum* L.). Corn stover is currently the feedstock of choice due to its current widespread availability and economic potential [10, 2]. Switchgrass is a promising perennial bioenergy crop that can be grown on marginal lands [11] and provides superior environmental benefits compared to corn, including support for biological diversity [12], lower nitrous oxide emissions [13], and improved soil properties [14, 15]. In order to investigate how interannual variation in precipitation influences the processing characteristics and microbial fermentation of these two important biofuel feedstocks, we compared switchgrass and corn stover that were harvested following the 2012 Midwestern U.S. drought to those harvested during two non-drought years with different precipitation patterns (2010 and 2013). In order to generate fermentable sugars, these materials were processed using ammonia fiber expansion (AFEX) pretreatment followed by enzymatic hydrolysis. The chemical composition of the feedstocks and hydrolysates were analyzed and the hydrolysates were fermented separately by *Saccharomyces cerevisiae* and *Zymomonas mobilis*. We also used a chemical genomics approach to evaluate the yeast biological response to the different hydrolysates.

## Results

### Drought year switchgrass hydrolysate is inhibitory to *Saccharomyces cerevisiae* growth and fermentation

Corn stover (Pioneer 35H56 and P0448R) and switchgrass (Shawnee and Cave-in-Rock) were harvested from the Arlington Agricultural Research Station (ARL) in south central Wisconsin from three growing seasons (2010, 2012, and 2013) that represent, with respect to total precipitation, an average year (2010), a major drought year (2012), and a year that was wet during the first half of the growing season and dry during the second half (2013) (Fig. 1). Each feedstock was processed using AFEX pretreatment and subjected to high solid loading enzymatic hydrolysis [6% and 7% glucan-loading for AFEX-treated corn stover hydrolysates (ACSH) and AFEX-treated switchgrass hydrolysates (ASGH), respectively] at previously optimized conditions [16]. Engineered xylose-utilizing ethanologens, *S. cerevisiae* Y128 [17] and *Z. mobilis* 2032 [18], were used to compare cell growth, glucose and xylose utilization, and ethanol production in the hydrolysates produced from corn stover and switchgrass harvested in different years. *Z. mobilis* exhibited similar growth, sugar utilization, and ethanol production for all hydrolysates, with slightly lower final cell densities but greater xylose consumption in the switchgrass hydrolysates (Fig. 2, Table 1). *S. cerevisiae* showed similar growth in all corn stover hydrolysates, but reduced xylose consumption in drought-year 2012 ASCH (P0448R) (Fig. 3A-D, Table 1). In the 2010 and 2013 ASGH, *S. cerevisiae* grew and consumed xylose more slowly than in the corn stover hydrolysates harvested in the same years (Fig. 3E,G, Table 1), but completely failed to grow or ferment glucose or xylose in the drought-year 2012 ASGH (Fig. 3F). With the exception of the *S. cerevisiae* fermentation of 2012 ASGH, all of the fermentations achieved final ethanol concentrations of between 30 − 40 g/L and ethanol yields of between ~200 ‒ 300 L/Mg untreated dry biomass (~45-70% of theoretical maximum) (Table 1).

**Fig. 1:**
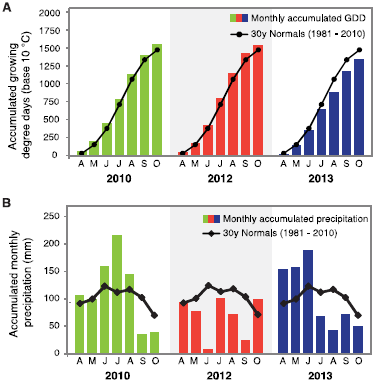
Interannual weather variation. **A)** Temperature (growing degree days (GDD)) and **B)** precipitation for 2010, 2012 and 2013, and the 30 year average values at Arlington Research Station in south-central Wisconsin (ARL, 43˚17’45" N, 89˚22’48" W, 315 masl).

**Fig. 2:**
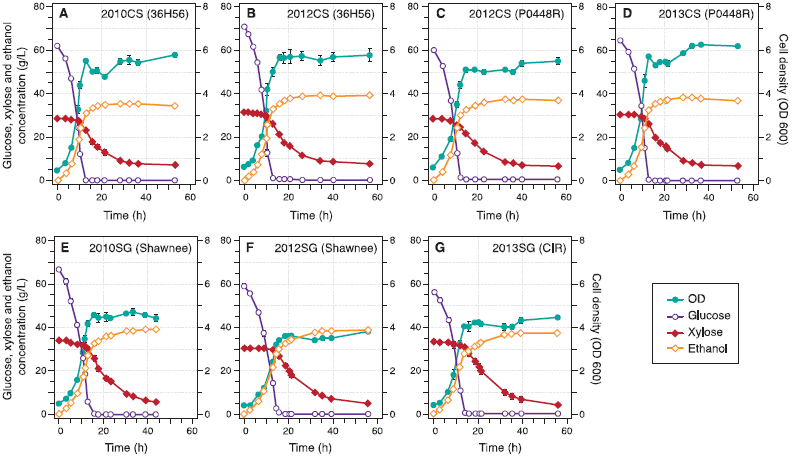
Fermentation profiles for *Zymomonas mobilis* 2032 grown in corn stover and switchgrass hydrolysates from different harvest years. **A)** 2010 ACSH (36H56), **B)** 2012 ACSH (36H56), **C)** 2012 ACSH (P0448R), **D)** 2013 ACSH (P0448R), **E)** 2010 ASGH (Shawnee), **F)** 2012 ASGH (Shawnee), **G)** 2013 ASGH (Cave-in-Rock (CIR)). Data points represent the mean ± s.d. (n=3). Error bars that are smaller than the individual data points may be hidden from view.

**Fig. 3:**
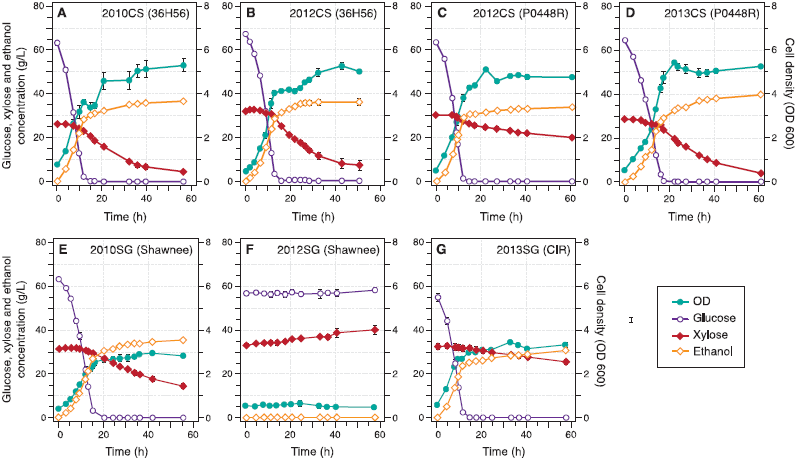
Fermentation profiles for *Saccharomyces cerevisiae* Y128 grown in AFEX-treated biomass hydrolysates. **A)** 2010 ACSH (36H56), **B)** 2012 ACSH (36H56), **C)** 2012 ACSH (P0448R), **D)** 2013 ACSH (P0448R), **E)** 2010 ASGH (Shawnee), **F)** 2012 ASGH (Shawnee), **G)** 2013 ASGH (Cave-in-Rock (CIR)). Data points represent the mean ± s.d. (n=3). Error bars that are smaller than the individual data points may be hidden from view.

**Table 1:** Summary of hydrolysis and fermentation results.

|  | Corn Stover |  |  |  | Switchgrass |  |  |
| --- | --- | --- | --- | --- | --- | --- | --- |
|  | 36H56 |  | P0448R |  | Shawnee |  | CIR |
|  | 2010 | 2012 | 2012 | 2013 | 2010 | 2012 | 2013 |
| Enzymatic Hydrolysis |  |  |  |  |  |  |  |
| Glucose Conversion (%)<sup>*</sup> | 92.7 ± 2.0 | 92.6 ± 1.5 | 94.4 ± 1.4 | 99.0 ± 3.0 | 74.4 ± 0.9 | 69.6 ± 2.4 | 75.1 ± 1.0 |
| Xylose Conversion (%)<sup>†</sup> | 67.3 ± 0.9 | 69.3 ± 1.2 | 67.0 ± 4.0 | 77.1 ± 4.2 | 63.4 ± 0.8 | 61.7 ± 5.0 | 69.6 ± 2.7 |
| Glucose Concentration (g/L) | 64.1 ± 1.4 | 64.0 ± 1.0 | 66.5 ± 1.0 | 67.6 ± 2.0 | 59.2 ± 0.8 | 60.3 ± 2.1 | 59.4 ± 0.8 |
| Xylose Concentration (g/L) | 27.3 ± 0.4 | 31.7 ± 0.5 | 28.9 ± 1.7 | 30.6 ± 1.7 | 31.2 ± 0.4 | 30.7 ± 2.5 | 35.3 ± 1.4 |
| Fermentation: *Zymomonas mobilis* |  |  |  |  |  |  |  |
| Final Time (hr) | 53.0 | 56.5 | 56.3 | 53.0 | 43.7 | 55.8 | 55.8 |
| Final Xylose Concentration (g/L) | 7.0 ± 1.1 | 7.7 ± 1.2 | 6.6 ± 0.4 | 6.5 ± 0.6 | 5.5 ± 0.3 | 4.9 ± 0.4 | 4.1 ± 0.6 |
| Final Ethanol Concentration (g/L) | 34.6 ± 0.6 | 39.4 ± 0.4 | 36.9 ± 0.7 | 36.7 ± 0.8 | 39.0 ± 1.2 | 38.9 ± 0.8 | 37.3 ± 0.5 |
| Metabolic Yield (%)<sup>‡</sup> | 80.4 ± 2.8 | 87.9 ± 2.1 | 81.4 ± 2.9 | 78.5 ± 3.6 | 90.2 ± 3.1 | 88.7 ± 4.3 | 80.6 ± 2.3 |
| Process Yield (%)<sup>§</sup> | 74.2 ± 2.4 | 80.8 ± 1.5 | 75.7 ± 2.8 | 73.3 ± 3.4 | 84.6 ± 3.1 | 83.9 ± 4.1 | 77.1 ± 2.1 |
| Ethanol Yield (L/Mg Untreated Dry Biomass) | 230 ± 4 | 262 ± 2 | 255 ± 5 | 288 ± 6 | 246 ± 7 | 214 ± 5 | 245 ± 3 |
| Max Theoretical Ethanol Yield (L/Mg Untreated Dry Biomass)<sup>¶</sup> | 371 ± 2 | 389 ± 2 | 401 ± 2 | 432 ± 1 | 415 ± 1 | 382 ± 3 | 437 ± 2 |
| Ethanol Yield (% of Maximum) | 61.8 ± 1.8 | 67.3 ± 1.1 | 63.6 ± 1.9 | 66.7 ± 2.1 | 59.4 ± 3.0 | 56.0 ± 2.3 | 56.2 ± 1.4 |
| Fermentation: *Saccharomyces cerevisiae* |  |  |  |  |  |  |  |
| Final Time (hr) | 56.5 | 50.8 | 60.8 | 60.8 | 55.5 | 57.3 | 57.3 |
| Final Xylose Concentration (g/L) | 4.7 ± 1.0 | 7.3 ± 4.5 | 19.9 ± 1.7 | 4.0 ± 0.6 | 14.2 ± 2.1 | 40.1 ± 3.7 | 25.4 ± 2.0 |
| Final Ethanol Concentration (g/L) | 36.6 ± 0.9 | 36.3 ± 2.9 | 34.0 ± 1.0 | 39.8 ± 0.3 | 35.3 ± 0.5 | 0.2 ± 0.0 | 30.6 ± 0.5 |
| Metabolic Yield (%)<sup>‡</sup> | 82.8 ± 3.1 | 80.5 ± 9.5 | 88.4 ± 4.5 | 82.9 ± 2.9 | 90.7 ± 3.3 | -5.4 ± 75.2 | 86.5 ± 4.0 |
| Process Yield (%)<sup>§</sup> | 78.6 ± 2.9 | 74.3 ± 8.0 | 69.9 ± 3.6 | 79.4 ± 2.8 | 76.5 ± 1.6 | 0.4 ± 11.6 | 63.3 ± 2.3 |
| Ethanol Yield (L/Mg Untreated Dry Biomass) | 243 ± 6 | 241 ± 19 | 236 ± 7 | 312 ± 2 | 223 ± 3 | 1 ± 0 | 202 ± 3 |
| Max Theoretical Ethanol Yield (L/Mg Untreated Dry Biomass)<sup>¶</sup> | 371 ± 2 | 389 ± 2 | 401 ± 2 | 432 ± 1 | 415 ± 1 | 382 ± 3 | 437 ± 2 |
| Ethanol Yield (% of Maximum) | 65.5 ± 2.4 | 61.9 ± 8.0 | 58.7 ± 2.9 | 72.2 ± 0.7 | 53.7 ± 1.4 | 0.3 ± 11.1 | 46.2 ± 1.6 |

Values are reported as the mean ± s.d. (n=3). Propagation of error was conducted to obtain s.d. values for all calculated values.
<sup>\*</sup>The glucose conversion was calculated based on the total glucan, soluble glucose, and glucose contributed by sucrose in the untreated biomass using a previously reported equation [51].
<sup>†</sup>The xylose conversion was calculated as: $Xyl/[BL \cdot Xln(150/132)]$ , where Xyl = hydrolysate xylose concentration (g/L), BL = biomass loading (g/L), Xln = untreated biomass xylan content (g/g biomass), and 150/132 are the molecular weights of xylose/xylan.
<sup>‡</sup>The metabolic yield is the ratio of sugars (glucose and xylose) consumed during fermentation to ethanol produced assuming 0.51 g ethanol/g sugars as the theoretical maximum.
<sup>§</sup>The process yield is the ratio of sugars initially present in the hydrolysate (glucose and xylose) to ethanol produced assuming 0.51 g ethanol/g sugars as the theoretical maximum.
<sup>¶</sup>The maximum theoretical ethanol yield is calculated based on the complete conversion of all glucose (as glucan, free glucose, or part of sucrose) and xylose (as xylan) in the untreated biomass to ethanol assuming 0.51 g ethanol/g sugars.

### Chemical genomic analysis of hydrolysates reveals a distinct pattern for drought year switchgrass hydrolysate

Chemical genomic analysis was used to measure the relative fitness of ~3500 single-gene deletion yeast strains [19] in the hydrolysates compared to synthetic hydrolysate [16] (**SI Dataset**). This analysis revealed a growth sensitivity profile of the 2012 ASGH that was drastically different from all other tested hydrolysates (Fig. 4A), which displayed profiles similar to those seen for ACSH and ASGH in a previous study [16]. The two most resistant mutants to the 2012 ASGH are *kex2*□, and *vps5*□, (Fig. 4B), the first of which encodes a protein residing in the trans-Golgi network [20], and the latter is part of the retromer complex for recycling of proteins from the late endosome to the Golgi apparatus [21]. Of the mutants that were highly susceptible in at least one of the hydrolysates (fitness < −2.5), 65 (16%) were susceptible to all five hydrolysates (Fig. 4C), with enrichment (p<0.05) in genes related to amino acid biosynthesis (**Fig. S1**). In contrast, of the 224 mutants that were highly resistant in at least one of the hydrolysates, only three were highly resistant to all five hydrolysates (fitness > 2.5) (Fig. 4C): *ygr237c*□, *ydr474c*□, and *bckl*□. The contrast between the 2012 ASGH and the other four feedstocks is reflected in the fact that 57 (14%) and 42 (19%) of highly susceptible and resistant mutants, respectively, were only highly susceptible or resistant to the 2012 ASGH (Fig. 4C). When the highly resistant mutants were limited to only those that had a statistically significant difference compared to the other four hydrolysates (p<0.001, n=42), gene ontology (GO) terms were enriched (p<0.05) for mutations related to Golgi/vesicle-mediated/vacuolar/endosomal transport and ribosome subunits (**Fig. S2A**). The next largest intersection was for mutants that were highly resistant or susceptible to all hydrolysates except the 2012 ASGH. When limited to highly susceptible mutants that had a statistically significant difference for the four hydrolysates compared to the 2012 ASGH (n=56, p<0.001), GO terms were enriched (p<0.05) related to the mitochondrial-nucleus signaling pathway, and Golgi/vacuolar transport (**Fig. S2B**). No significant terms were found for the mutants that were only highly susceptible to the 2012 ASGH or only highly resistant to the other four feedstocks. Gene set enrichment analysis was used to evaluate whether any yeast metabolic pathways (using the KEGG pathway collection) were enriched for the mutants that were significantly different between the 2012 ASGH and the four other feedstocks (p<0.001). This analysis revealed three KEGG pathways (FDR<0.25) that were dominated by mutants that were resistant to the 2012 ASGH and susceptible to the hydrolysates of the other four feedstocks: SNARE interactions in vesicular transport, endocytosis, and the ribosome. For the SNARE pathway, the gene deletions that conferred greater resistance in 2012 ASGH compared to the other hydrolysates (p<0.001) were *GOS1*, *VAM7*, and *SEC22*, which are all involved in vesicle traffic between the ER, Golgi, endosome, and vacuole.

**Fig. 4:**
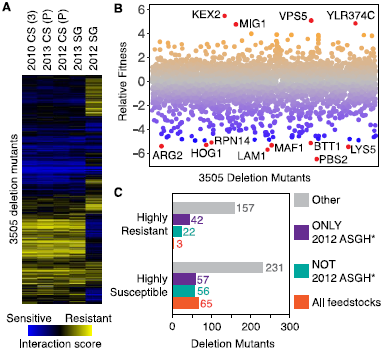
Chemical genomic analysis of hydrolysate variation. **A)** Fitness heat map for yeast mutants in corn stover (CS) and switchgrass (SG) hydrolysates. The genome-wide yeast deletion mutant collection was grown in fifteen different hydrolysate batches (n=3 per feedstock) and a synthetic hydrolysate (SynH2.1) control (n=3). The abundance of each mutant was assessed by sequencing the strain specific barcodes and a fitness score was determined relative to the synthetic hydrolysate control. Mutants sensitive to the hydrolysate conditions are shown in blue and resistant are shown in yellow, compared to the abundance in the SynH2.1 control. The (3) represents the 36H56 variety and the (P) represents the P0448R variety of corn stover. **B)** Fitness plot of yeast mutants grown in 2012 ASGH. The most resistant (fitness >4) and susceptible mutants (fitness <−5) are labeled and shown in red. **C)** Intersection of yeast mutants that are highly susceptible or resistant to all hydrolysates, only the 2012 ASGH, or all hydrolysates except the 2012 ASGH. ^*^The fitness of these mutants was statistically different (p<0.001) in the 2012 ASGH versus the other four hydrolysates [2013 ASGH, 2010 CS (36H56), 2012 CS (P0448R), 2013 CS (P0448R)].

### Imidazoles and pyrazines are present in high concentrations in drought year switchgrass hydrolysate

To identify the cause of severe growth inhibition in the 2012 ASGH, we compared the compositions of the untreated biomass (Fig 5, **Table S1**), hydrolysates (**Tables S2-S4**) and extracts of the pretreated biomass. As is typical for drought stressed grasses [7, 22], untreated 2012 switchgrass contained higher total extractives (water- and ethanol-extractable compounds) and soluble sugars (Fig. 5A) and lower structural carbohydrates and lignin compared to the 2010 and 2013 switchgrass (Fig. 5B). A number of amino acids, metals, and furanic and phenolic compounds were also directly quantified from the hydrolysates (**Tables S2-S4**). With the exception of the 2010 and 2013 ASGH, which overlapped, all the hydrolysates were readily distinguishable by principal component analysis (PCA) of their hydrolysate compositions (Fig. 6). The greatest amount of variation (31%) was attributed to the difference between plant species (corn stover vs. switchgrass) (Fig. 6A), followed by the difference between 2010/2013 and 2012 hydrolysates (22% of variance) (Fig. 6B). Of all the compounds in the hydrolysate, the amino acid content had the largest influence on segregation of the 2012 feedstocks (Fig. 6C). When looking at the compounds individually, compared to the other hydrolysates, the 2012 ASGH had statistically higher (p<0.05) levels of benzamide (10 μM), vanillyl alcohol (0.8 μM), sulfur (5.4 mM), chloride (96.6 mM – largely from HCl used to neutralize the hydrolysate), magnesium (24.4 mM), total nitrogen (307.3 mM), proline (1.46 mM), and tryptophan (42.5 μM).

**Fig. 5:**
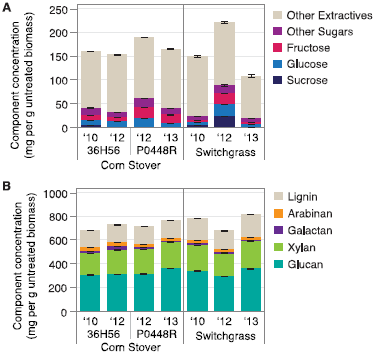
Untreated biomass composition. **A)** Water and ethanol soluble extractives. **B)** Structural carbohydrates and lignin. Values are reported as the mean ± s.d. (n=3).

**Fig. 6:**
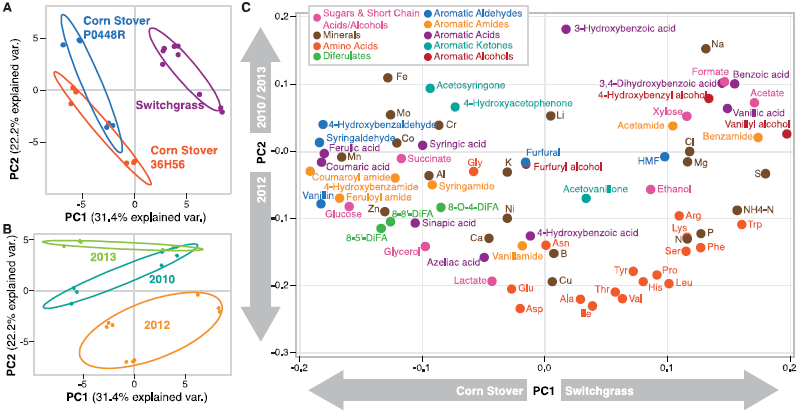
Principal component analysis (PCA) of hydrolysate composition data – Relationship between principal component 1 and 2. **A)** Hydrolysate batches grouped by plant variety. **B)** Hydrolysate batches grouped by year. **C)** Correlation score graph showing relative effect of each hydrolysate component.

In order to determine whether any additional compounds were present that might be responsible for the inhibition, the hydrolysates were extracted with ethyl acetate and analyzed. These extracts revealed the presence of higher levels of pyrazines in the drought-year (2012) ASGH compared to the other switchgrass hydrolysates (Fig. 7A). Seven substituted imidazoles and pyrazines were further quantified from acetone extracts of the untreated and pretreated biomass. These compounds were found at higher levels in pretreated biomass samples and were either present at very low concentrations (imidazoles) or absent in the untreated biomass, indicating that they were produced during the AFEX pretreatment process (Fig. 7B). Pretreated switchgrass contained more pyrazines than pretreated corn stover, and the drought-year (2012) switchgrass exhibited the highest concentration of pyrazines. Combined imidazole and pyrazine levels after pretreatment correlated with the soluble sugar content of the untreated biomass (Fig. 7C). The concentrations of imidazoles and pyrazines in the hydrolysates were estimated based on their concentrations in the pretreated biomass (Table 2). The total estimated concentration of all imidazoles and pyrazines in the 2012 ASGH was almost twice that of the next highest sample, 2013 ACSH (P0448R) (Table 2), and the concentrations of 2-methylimidazole, 4(5)-methylimidazole, and 2-methylpyrazine were higher than the majority of the other aromatic compounds that were characterized in the 2012 ASGH (Table 2, **Table S2**). Acetamide and four of the top five most abundant low molecular weight phenolics (coumaroyl amide, feruloyl amide, coumaric acid, and vanillin) were at higher levels in the readily fermentable 2012 ACSH (36H56) compared to the inhibitory 2012 ASGH (Table 2).

**Fig. 7:**
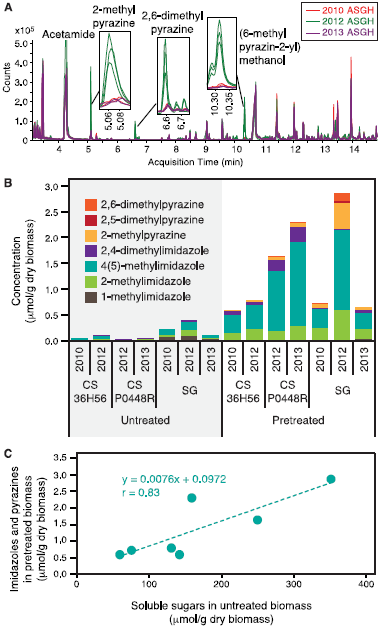
Imidazole and pyrazine detection and quantification in AFEX-treated biomass and hydrolysates. **A)** Overlaid mass spectrometric chromatogram of ethyl acetate extracts of AFEX-treated switchgrass hydrolysates. Each line represents a replicate batch of hydrolysate (2012: n=3; 2010 and 2013: n=2). **B)** Imidazole and pyrazine content of untreated and AFEX-treated corn stover (CS) and switchgrass (SG). **C)** Correlation between imidazole and pyrazine content of AFEX-treated biomass and untreated biomass soluble sugars (sucrose, glucose fructose, xylose, arabinose, and galactose).

**Table 2:** Concentrations (μM) of imidazoles and pyrazines (estimated) and aromatic degradation products in pretreated biomass hydrolysates

|  | Corn Stover 36H56 |  | Corn Stover P0448R |  | Switchgrass |  |  |
| --- | --- | --- | --- | --- | --- | --- | --- |
|  | 2010 | 2012 | 2012 | 2013 | 2010 | 2012 | 2013 |
| 1-methylimidazole | 2 | 0 | ND | 0 | 1 | 1 | 5 |
| 2-methylimidazole | 26 | 42 | 33 | 44 | 45 | 137 | 37 |
| 4(5)-methylimidazole | 67 | 86 | 214 | 264 | 76 | 356 | 60 |
| 2,4-dimethylimidazole | 13 | 14 | 36 | 49 | 5 | 6 | 12 |
| 2-methylpyrazine | 4 | 6 | 11 | 10 | 14 | 114 | 10 |
| 2,5-dimethylpyrazine | 0 | 0 | 1 | 1 | 1 | 9 | 1 |
| 2,6-dimethylpyrazine | 1 | 1 | 4 | 4 | 3 | 38 | 2 |
| Sum of imidazoles & pyrazines | 112 | 150 | 299 | 371 | 144 | 661 | 126 |
| Acetamide | 7755 | 7114 | 8283 | 8309 | 8888 | 11372 | 8981 |
| Coumaroyl amide | 5034 | 3152 | 3763 | 4502 | 1622 | 1751 | 1908 |
| Feruloyl amide | 2044 | 1529 | 2181 | 2186 | 593 | 1179 | 753 |
| Coumaric acid | 1328 | 489 | 774 | 937 | 225 | 268 | 257 |
| Benzoic acid | 132 | 168 | 121 | 124 | 304 | 225 | 300 |
| Vanillin | 181 | 141 | 156 | 146 | 71 | 69 | 86 |

### Imidazoles and pyrazines contribute to the inhibition of *S. cerevisiae*

In order to determine whether elevated imidazoles and pyrazines contribute to the anaerobic growth inhibition of *S. cerevisiae* Y128 in the 2012 ASGH, we added these compounds into the non-inhibitory 2010 ASGH at the levels estimated in 2012 ASGH and up to 50 times the concentration. Prior to supplementation with additional imidazoles and pyrazines, the 2010 ASGH supported yeast growth and fermentation (Fig. 3E, Fig. 8). While there was still growth at the concentration of imidazoles and pyrazines in the 2012 ASGH (1X), growth began to be delayed at 25 times the concentration (25X), with complete inhibition at 50 times the concentration (50X) within the fermentation time frame. These results correspond to the IC_50_ values, where at comparable concentrations the individual imidazoles and pyrazines reduced growth of *S. cerevisiae* by 50%, with the imidazoles more strongly inhibitory (Table 3).

**Fig. 8:**
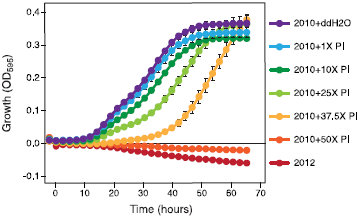
Imidazoles and pyrazines found in drought year AFEX-treated switchgrass hydrolysate (ASGH) can impair anaerobic yeast growth. Anaerobic yeast growth in add-back experiment, with various concentrations of pyrazines and imidazoles (P/I) in 2010 ASGH relative to estimated levels in 2012 ASGH (mean, n=3). Average cell densities with standard error of the mean are reported from triplicate samples, with every twelfth time point plotted (roughly one time point every two hours).

**Table 3:**
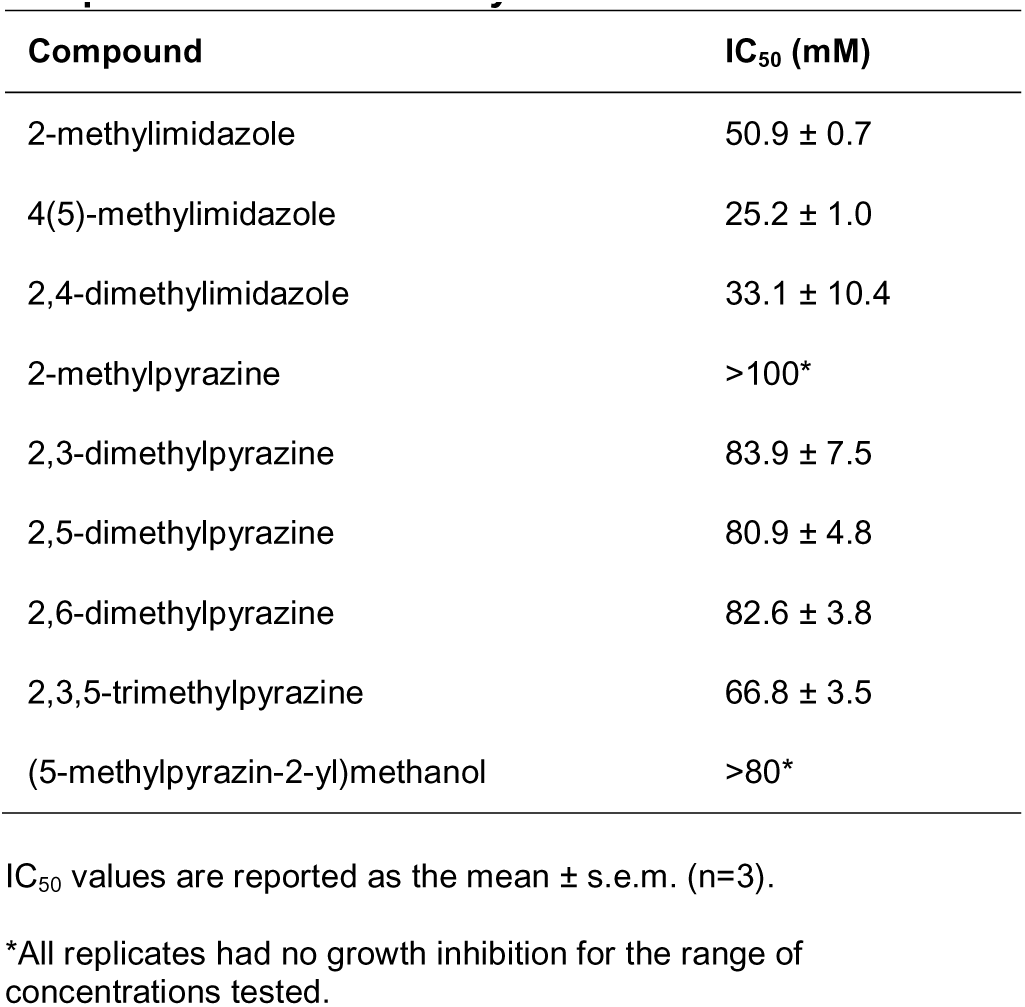
IC_50_values of selected nitrogenous compounds for *Saccharomyces cerevisiae* Y128

| Compound | IC<sub>50</sub> (mM) |
| --- | --- |
| 2-methylimidazole | 50.9 ± 0.7 |
| 4(5)-methylimidazole | 25.2 ± 1.0 |
| 2,4-dimethylimidazole | 33.1 ± 10.4 |
| 2-methylpyrazine | >100* |
| 2,3-dimethylpyrazine | 83.9 ± 7.5 |
| 2,5-dimethylpyrazine | 80.9 ± 4.8 |
| 2,6-dimethylpyrazine | 82.6 ± 3.8 |
| 2,3,5-trimethylpyrazine | 66.8 ± 3.5 |
| (5-methylpyrazin-2-yl)methanol | >80* |

IC<sub>50</sub> values are reported as the mean ± s.e.m. (n=3).
\*All replicates had no growth inhibition for the range of concentrations tested.

## Discussion

During the severe Midwestern drought in 2012, soluble sugars accumulated to significantly higher levels in switchgrass compared to during two non-drought years in 2010 and 2013. During ammonia-based pretreatment (AFEX), these soluble sugars underwent Maillard reactions with ammonia to form aromatic nitrogenous compounds, imidazoles and pyrazines [23, 24]. Both classes of compounds can be highly toxic [26] and many complex azoles are potent antifungal agents [25]. Our data suggest that these compounds contributed to inhibition of fermentative yeast growth in drought stressed switchgrass (Fig. 9), however they are most likely not the sole cause. A previous study predicted reductions of 10-15% in the theoretical ethanol yield from lignocellulosic biomass harvested during a drought year compared to a non-drought year, largely due to the negative effects of drought on the biomass structural sugar content [7]. In our study, while in some cases there was a reduction in the actual ethanol yield for drought year feedstocks (−7% for CS-P0448R and SG), this was not always the case (+12% for CS-36H56 for 2012 vs. 2010) (Table 1). The actual ethanol yield also varied significantly between feedstocks (from 46 – 72% of the theoretical maximum) in a manner that was not obvious based on the untreated biomass composition. Additionally, the complete inhibition of the yeast hydrolysate in the 2012 ASGH, while related to the biomass composition, was not predictable based on the current state of knowledge. In order to design feedstocks and processes that are able to either accommodate or reduce feedstock variability, more studies are needed that focus on understanding how external factors influence biomass quality and subsequently affect fermentation performance.

**Fig. 9:**
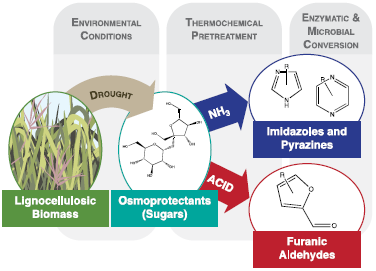
Interaction between plant response toenvironmental conditions and pretreatment chemistry. Inlignocellulosic biomass, drought stress causes an increase in osmoprotectants, including soluble sugars that are degraded to microbial inhibitors during thermochemical pretreatments.

Although the drought had some negative effects on hydrolysate composition, it also had a number of positive effects, particularly related to hydrolysate amino acid concentrations. With the exception of glycine and asparagine, the drought year hydrolysate for each respective feedstock, had the highest concentration of each amino acid, and of all hydrolysates the 2012 ASGH had the highest concentration for both proline and tryptophan (**Table S4**). Plants commonly respond to drought or other abiotic stresses by accumulating amino acids [26, 5]. In particular, proline is produced by drought stressed plants to help regulate osmotic pressure [5] and both proline and tryptophan have been reported at higher levels in drought stressed grasses compared to their unstressed counterparts [6, 22]. For pretreatments, such as AFEX, that do not denature, degrade, or remove proteins and amino acids, the retention of amino acids in the hydrolysate provides a beneficial source of nutrients for the microorganism [27]. The importance of these amino acids to microbial fitness in the hydrolysates is apparent from the large number of amino acid biosynthetic mutants that were highly susceptible in all of the five hydrolysates investigated (**Fig. S1**).

In our study, the soluble sugars that were present in the lignocellulosic biomass were degraded to inhibitory imidazoles and pyrazines following ammonia-based pretreatment. However, for other pretreatment methods, the soluble sugars that accumulate in drought stressed biomass can also be degraded to other inhibitory compounds, in the case of dilute acid to furfural, 5- hydroxymethylfurfural, levulinic acid and formic acid [28]. These compounds can cause severe negative effects on microbial fermentation for both yeast and bacteria [29, 30]. Thus, degradation of soluble sugars that are present in drought stressed crops poses a potential problem for lignocellulosic biofuel production regardless of the pretreatment used. However, it may be possible to overcome the inhibition by adjusting pretreatment conditions to limit formation of harmful compounds, removing soluble sugars prior to processing, or utilizing more resistant microbial strains. For example, it may be preferable to use the bacterium *Z. mobilis* 2032, which was less susceptible to growth inhibition in the 2012 ASGH compared to the yeast *S. cerevisiae* Y128 (Fig. 2–3).

Analysis of the chemical genomics data indicate that the 2012 ASGH had an impact on the protein trafficking system within the yeast cell, particularly in relationship to the late endosome and retromer, which is responsible for recycling of certain proteins from the late endosome to the Golgi apparatus. In yeast the retromer consists of two subcomplexes, a trimer consisting of Vps26p, Vps29p, Vps35p and a dimer consisting of Vps5p and Vps17p [21]. A number of mutants related to these systems, in particular the three retromer subunits for which we had mutants (*vps35*□, *vps5*□, and *vps17*□), were highly susceptible to reduced growth in the four other hydrolysates that were investigated (2010, 2012-P0448R and 2013 ACSH, and 2013 ASGH) but had greater fitness in the 2012 ASGH. If the mechanism of inhibition in the 2012 ASGH is related to the endosomal system and vesicular transport between the organelles, this could explain the difference observed with the bacterial ethanologen *Z. mobilis*, which has neither organelles nor the process of endocytosis, and was able to grow with no difficulty in the 2012 ASGH.

Plants experience drought stress in response to low levels of soil moisture. Although there are benefits to growing dedicated bioenergy crops like switchgrass on marginal lands to avoid competition with food crop production [31], some marginal lands are classified as such because their soil has poor water holding capacity [32]. Plants grown on these soils may experience greater drought stress and produce more osmoprotective soluble sugars than plants grown on more fertile soils. Climate change may further aggravate these issues as extreme precipitation events are predicted to increase [33]. Intense rainfall followed by longer dry spells limits the replenishment of soil moisture [33], and in certain regions this may negatively influence biomass yields and processing characteristics. Moisture stress will be an issue for bioenergy production systems that needs to be addressed, not just because of the impact on crop yields, but also because of the potential negative impact on biomass quality.

## Conclusions

Drought induces the accumulation of high concentrations of soluble sugars in lignocellulosic bioenergy crops. During ammonia-based pretreatment these sugars are degraded to imidazoles and pyrazines that during fermentation contribute to growth inhibition of the yeast *S. cerevisiae*, but do not negatively effect the bacterium *Z. mobilis*. This is the first study that links compounds generated during the processing of environmentally stressed lignocellulosic biomass to deleterious impacts on the microbes during biofuel production. Our findings have profound implications for the development of sustainable lignocellulosic biofuel production systems that are able to tolerate fluctuations in precipitation and water availability.

## Methods

The methods for AFEX pretreatment; high solids enzymatic hydrolysis; chemical analysis of hydrolysate composition; and strains, media, growth and fermentation conditions are the same as previously reported [16].

### Feedstock Production, Harvest, and Processing

Switchgrass and corn stover were cultivated at the Arlington Agricultural Research Station (ARL, 43^°^17’45" N, 89^°^22’48" W, 315 masl) in Arlington, Wisconsin. Corn stover was sourced from Arlington field 744 (ARL-744) in 2010, ARL-570 in 2012, and ARL-742 in 2013. Switchgrass was sourced from ARL-346 in both 2010 and 2012, and ARL-115 in 2013. The main soil at ARL is Plano silt-loam (fine-silty, mixed, superactive, mesic Typic Argiudoll); a deep (>1 m), well-drained mollisol developed over glacial till and formed under tallgrass prairie [13]. Mean annual temperature and precipitation are 6.9 ^°^C and 869 mm, respectively [34, 35].

Pioneer 36H56 and P0448R corn stover (both triple stacked with Roundup Ready and corn borer and rootworm resistance) were planted on May 3 (2010) and May 11 (2012) for 36H56, and May 11 (2012) and May 15 (2013) for P0448R. Fertilizer (0-0-50 potassium sulfate) was applied in 2010 after 4 years of alfalfa. In 2012, both corn varieties received 92 kg N ha^−1^ as anhydrous in April, whereas 2013 corn received 83 kg N ha^−1^ as urea in May. Weed control was attended on ARL-744 with a pre-emerge (Metolachlor: 1848 mL AI ha^−1^) and post-emerge herbicide (Dicamba; Diflufenzopyr: 267 mL AI ha^−1^) applied on May 10 and June 10 2010, respectively. For field ARL-570, a mixed pre-emerge herbicide (2,4-D LV4 Ester; Glyphosate; Mesotrione; S-Metolachlor: 1264 mL AI ha^−1^) was applied on April 16, 2012 prior to planting and a mixed post-emerge herbicide (Glyphosate; Tembotrione; Ammonium Sulfate; Methylated Seed Oil: 852 mL AI ha^−1^) on June 8, 2012. The herbicide treatment on ARL-742 used Mesotrione: 175 mL AI ha^−1^ + S-Metochlor: 1685 mL AI ha^−1^. Corn stover was collected shortly after grain harvest in early November of all years using a combine that had been modified to separate the corn grain and then chop and bail the corn stover.

Switchgrass (Shawnee variety; 2010 and 2012) was planted on May 29, 2004 using a Brillion Sure Stand seeder (Landoll Corporation, Marysville, KS) at a rate of 16.8 kg ha^−1^. For initial weed control, Quinclorac herbicide (1445 mL AI ha^−1^) was applied one day after planting. A tank mix of Imazethapyr (259 ml AI ha^−1^) and Dicamba (1445 mL AI ha^−1^) was applied on May 19, 2006 for additional weed control. Each year in April granular urea (46-0-0) was top-dressed at a rate of 90 kg ha^−1^. In mid-October 2010, switchgrass was cut and conditioned with a 4.5 m wide hay-bine (John Deere 4990). Switchgrass sourced in 2013 (Cave-in-Rock) was planted in late June 2008 using a drop spreader (Truax Company, Inc.) with two culti-pack rollers at a rate of 14 kg ha^−1^. Initial weed control was accomplished with Glyphosate (700 mL AI ha^−1^) on June 17, 2008 and again as a pre-emerge treatment on April 23, 2009, and May 3, 2010. Post-emerge weed control was applied as 2,4-D (773 mL AI ha^−1^) on June 26, 2009 and May 10, 2010. Starting in 2010, 56 kg ha^−1^ (34-0-0 ammonium nitrate) was applied annually, and in 2013 N was applied on May 30. In mid-to late-September (2010 and 2012) and mid-October (2013), biomass was cut and windrowed, then chopped with a self-propelled forage harvester into a dump wagon equipped with load cells.

Following harvest, each corn stover and switchgrass material was dried in a 60 °C oven until the dry weight was stable (~48 h), then milled using a 18-7-301 SchutteBuffalo hammer mill (SchutteBuffalo, Buffalo, NY) equipped with a 5 mm screen, and stored at room temperature in sealed bags until use.

### Chemical Genomic Analysis of Hydrolysates

Chemical genomic analysis of these hydrolysates was performed as described previously using a collection of ~3500 yeast deletion mutants [36, 19]. 200 μL cultures of the pooled collection of *S. cerevisiae* deletion mutants were grown anaerobically in the different versions of ACSH and ASGH, or yeast rich medium (YPD, 20 g/L peptone, 10g/L yeast extract, 20 g/L glucose), diluted 1:1 with sterile water, in triplicate for 48 h at 30 ^°^C. Genomic DNA was extracted from the cells and mutant-specific molecular barcodes were amplified using specially designed multiplex primers as described previously [19]. The barcodes were sequenced using an Illumina HiSeq2500 in rapid run mode (Illumina, Inc., San Diego, CA). The barcode counts for each yeast deletion mutant in the hydrolysates were normalized against the synthetic hydrolysate control (SynH2.1)[16] in order to define sensitivity or resistance of individual strains (chemical genetic interaction score). The pattern of genetic interaction scores for all mutant strains represents the chemical genomic profile or “biological fingerprint” of a sample [36, 19]. The clustergram of the chemical genomic profiles was created in Cluster 3.0 [37], and visualized in Treeview (v1.1.6r4) [38]. The p-value for the difference between 2012 ASGH and all other hydrolysates was calculated and Bonferroni corrected using the multtest package [39] in R-Studio^®^. A Bonferroni-corrected hypergeometric distribution test was used to search for significant enrichment of GO terms among sets of highly resistant mutants (fitness > 2.5, n = 224) and highly susceptible mutants (fitness < −2.5, n = 409) [40] using LAGO [41]. For the highly resistant and susceptible mutants for only 2012 ASGH or the four feedstocks but not 2012 ASGH, the GO terms were evaluated using only those terms that had statistically different fitness between the two groups (p<0.001). Gene set enrichment analysis (GSEA) [42] was used to compare the enrichment of the KEGG pathways for *Saccharomyces cerevisiae* between the 2012 ASGH and the four other feedstocks for genes that conferred statistically different fitness (p<0.001).

### Untreated Biomass Composition Analysis

The composition of the untreated biomass was analyzed based on the NREL standard procedures for biomass composition analysis [43–47], with the following deviations. Samples for composition analysis were milled through a 2 mm screen using a Foss Cyclotec™ mill (Eden Prairie, MN) and not sieved prior to analysis. The protein content was estimated by multiplying the nitrogen content as determined by a Skalar Primacs SN Total Nitrogen Analyzer (Breda, The Netherlands) by a conversion factor (6.25), which assumes that 16% of the protein is nitrogen. Although the preferred method is to calculate the protein content based on the amino acid profile of the biomass [48], this method is complex and so we chose to use an estimation. Water-soluble oligomeric sugars were determined by hydrolyzing the water extractives using sulfuric acid [49]. The hydrolyzed water extracts were then neutralized using calcium carbonate and both the hydrolyzed and non-hydrolyzed water extractives were run through an Aminex HPX-87P column (Bio-Rad, Hercules, CA) with attached guard columns [47] and analyzed for their sucrose, fructose, glucose, xylose, arabinose, galactose, and mannose concentrations based on calibration standards.

### Hydrolysate Amino Acid Composition

Prior to amino acid quantification, 50 μL aliquots of samples were spiked with stable isotope labeled internal standards for the 20 common proteinogenic amino acids (Sigma-Aldrich Cell Free Amino Acid Mixture - ^13^C,^15^N; P/N 767964-1EA) and processed by solid-phase extraction (Phenomenex Strata-X-C cartridges; P/N 8B-S029-HCH) to remove matrix interferents. SPE-processed samples underwent vacuum centrifugation before resuspension in 1 mL of Mobile Phase A. Samples were then analyzed via LC-MS/MS, based on the protocol from Gu et al. [48], with the following modifications: mobile phase A was 10 mM instead of 1 mM (to reduce column equilibration time) and an LC gradient of: 0.00-1.75 min (98% A); 1.76-8.00 min (linear ramp to 45% A); 8.01-9.00 min (10 % A); 9.01-13.00 min (98% A). Response factors were calculated based on the peak area of the selected multiple reaction monitoring (MRM) chromatograms for each compound relative to the area of the MRM peak for each amino acid’s stable isotope labeled internal standard.

### Statistical Analysis of Hydrolysate Composition

Statistical analysis of the hydrolysate composition was conducted in R-Studio^®^, version 0.98.1102 (Boston, MA). A linear model of each chemical component was developed based on the feedstock (corn stover or switchgrass), harvest year and their interaction, with variety nested within feedstock. The model was evaluated using Tukey’s HSD test based on 95% confidence intervals (Agricolae package, version 1.2-1 [49]). When a reported value was below the limit of quantitation (LOQ), the value was recalculated as LOQ/√2 [50]. These recalculated values were used to determine the mean, standard deviation, and statistical differences. The principal component analysis was conducted in R-Studio^®^ and plots were generated using the ggbiplot package.

### Quantification of Imidazoles and Pyrazines in AFEX-treated Biomass

AFEX-pretreated biomass was milled through a 2.0 mm screen using a Foss Cyclotec™ mill (Eden Prairie, MN). The milled biomass was extracted with acetone using an Accelerated Solvent Extractor (Dionex™ ASE 200, Thermo Scientific) and the following conditions: 5 min heat, 5 min static, 150% flush volume,120 sec purge, 2 cycles, 1500 psi, and 70 ^°^C. Standards for the analyzed compounds were prepared in pure acetone in concentrations ranging from 0.00128 to 20 mg/L. Internal standards of 4-methylimidazole-d_6_ (imidazole authentic standard) and 2-methylpyrazine-d_6_ (pyrazine authentic standard) were obtained from C/D/N Isotopes (Pointe-Claire, Quebec, Canada) and added to each sample, standard, and blank at a final concentration of 6 mg/L. Samples were directly analyzed via GC-MS, without derivatization, based on the protocol from Chundawat et al. [23], with the following modifications to the GC temperature program: 40 °C (2 min), from 5 °C/min to 150 °C (1 min hold), 8 °C/min to 200 °C (2 min hold), 20 °C/min to 260 °C (3 min hold). Response factors were calculated based on the peak area of the selected ion chromatogram (molecular ion; M+) of each compound relative to the area of the internal standard peak.

### Determination of Pyrazines and Imidazoles in Switchgrass Hydrolysates by RP-HPLC-HR/AM-MS

For each year of ASGH, 1 mL of hydrolysate was extracted with 0.5 mL ethyl acetate by vortex mixing for approximately 30 seconds, then centrifuging at 16xg for 5 min to separate the layers. The organic (top) phase was collected and the procedure was repeated with another 0.5 mL ethyl acetate and the second organic extract combined with the first. 0.5 g anhydrous sodium sulfate was added to the ethyl acetate extract, capped, and allowed to stand overnight before an aliquot was taken for analysis by GC-MS. Sample components were separated by an Agilent 7890A Gas Chromatograph equipped with an HP-5 MS column, 30m x 0.25mm ID, initially at 40^°^C for 2 minutes then heated to 320 ^°^C at 10 ^°^C/min. Mass spectra were recorded with the interfaced Agilent 5975 MSD from m/z 40 to 750 with an ionization energy of 70eV. The GC inlet was set to 265 °C and MS transfer line temperature was 250 °C. The inlet was operated in spilt mode with a split ratio of 10:1, helium carrier gas flow rate through the column was held at 1 mL/min. Mass Hunter GC/MS acquisition software (Agilent) version B.07.00.1413 was used to control the instrument and collect the data. Identities of 2-methylpyrazine, 2,6-dimethylpyrazine, 2,3-dimethylpyrazine (the smaller peak immediately following the 2,6 isomer), (5-methylpyrazin-2-yl) methanol, and (6-methylpyrazin-2-yl) methanol were confirmed by co-chromatography with authentic reference standards. Known amounts of authentic reference standards were individually added to aliquots of a composite mixture of the ethyl acetate extracts of all three batches of 2012 ASGH. The resulting chromatograms were compared to the chromatogram of the composite mixture without added standards. Single point calibration gave a value of 300 μM 2-methylpyrazine in the extract. Further experiments suggested the actual value was higher due to incomplete extraction.

### Pyrazine and Imidazole Spike-In Experiment

To determine if the detected pyrazines and imidazoles contributed toward the inhibition of yeast in the 2012 ASGH, we grew yeast in 2010 ASGH supplemented with similar levels of pyrazines and imidazoles found in the 2012 ASGH. 10 μL of exponentially growing Y128 *S. cerevisiae* [17] at a cell density of OD_600_=1.0 was inoculated in 96-well microtiter plates containing 190 μL. We grew triplicate, 200 μL cultures of *S. cerevisiae* Y128 aerobically in each of the following hydrolysates: 2012 ASGH and 2010 ASGH supplemented with 0X, 1X, 10X, 25X, 37.5X, or 50X concentrations of the following pyrazines and imidazoles from 2012 ASGH dissolved in ddH_2_O:1 μM 1-methylimidazole, 140 μM 2-methylimidazole, 360 μM 4-methylimidazole, 6 μM 2,4-dimethylimidazole, 430 μM 2-methylpyrazine, 9 μM 2,5-dimethylpyrazine, 38 μM 2,6-dimethylpyrazine (Sigma, USA). Cultures were incubated at 30°C for 72 h and read every 11.3 minutes using a TECAN M1000 multimode plate reader housed within in an anaerobic chamber (Coy) maintained with 10% H_2_, 10% CO_2_ and 80% N_2_ gases.

### IC_50_ of Imidazoles and Pyrazines

To determine the half maximal inhibitory concentration (IC_50_) of each pyrazine and imidazole, we created a 12-point dose curve of each compound. We grew 200 μL cultures of *S. cerevisiae* Y128 in synthetic hydrolysate (SynH2.1)[16] supplemented with a range of 0 to 10 mg/mL (0, 0.5, 1, 2, 3, 4, etc.) of each of the following compounds separately: 2-methylimidazole, 4(5)-methylimidazole, 2,4-dimethylimidazole, 2-methylpyrazine, 2,3-dimethylpyrazine, 2,5-dimethylpyrazine, 2,6-dimethylpyrazine, 2,3,5-trimethylpyrazine, and (5-methylpyrazin-2-yl) methanol. We incubated these cultures for 48 hours with OD_595_ readings taken every 15 minutes using a TECAN M500 (TECAN, USA). Biological replicates were conducted in triplicate. IC_50_ values were estimated using SigmaPlot 12.0 (Systat Software, San Jose, CA) and converted to molar concentrations for ease of comparison.

#### List of Abbreviations

ACSH: AFEX-treated corn stover hydrolysate
ASGH: AFEX-treated switchgrass hydrolysate
AFEX: ammonia fiber expansion pretreatment
CS: corn stover
SG: switchgrass

## Declarations

### Ethics Approval and Consent to Participate

Not applicable

### Consent for Publication

Not applicable

### Availability of Data and Materials

Correspondence and reasonable requests for data or materials should be addressed to R.G.O.

## Author Contributions

RGO designed the project, performed composition analysis and prep work for pyrazine and imidazole quantification, analyzed data, ran statistical analyses, analyzed chemical genomics data, and wrote the manuscript with input from all authors. LGO, DE, and GRS designed the project, coordinated collection and processing of biomass, analyzed data, and edited the manuscript. YZ designed the project, led hydrolysate production and fermentation experiments, analyzed data, and edited the manuscript. JS, DX, and EP generated hydrolysate, conducted fermentation experiments, analyzed data, and edited the manuscript. SAS, AH, ADJ and JJC designed and performed mass spectrometric analysis of biomass extracts and hydrolysates, analyzed data, and edited the manuscript. JSP designed the project, conducted chemical genomics experiments, analyzed data and edited the manuscript. TKS, SB, and QD conducted chemical genomic experiments, individual compound toxicity tests, and spike in experiments, analyzed data, and edited the manuscript. DMB and DC designed the project, analyzed data, and edited the manuscript. All authors read and approved the final manuscript.

## Acknowledgements

We thank K. Keegstra, B. Landick, R. Jackson, J. Ralph, and B. Dale for feedback during preparation of the manuscript. We also thank Novozymes for providing the enzymes used during enzymatic hydrolysis; J. Sustachek, A. Miller, Z. Andersen, B. Faust, and J. Tesmer for collection and processing of biomass; C. Donald Jr. for AFEX pretreatment; M. Kreuger, M. Shabani, and C. Gunawan for biomass composition analysis; M. Kreuger for prep work for imidazole/pyrazine quantification; M. McGee for HPLC analysis of fermentation products; and the MSU Mass Spectrometry and Metabolomics Core for mass spectrometric analysis of biomass extracts and hydrolysates.

## Supporting Information

Supporting_Information_1.pdf: Maps of significant gene ontology terms for chemical genomics data. Untreated biomass composition. Detailed hydrolysate composition.

Supporting_Information_2.csv: Chemical genomics dataset.

